# Force Reserve Predicts Compensation in Reaching Movement with Induced Shoulder Strength Deficit

**DOI:** 10.1101/2024.03.28.587142

**Authors:** Germain Faity, Victor R. Barradas, Nicolas Schweighofer, Denis Mottet

**Author notes:** Correspondence: Denis Mottet.

## Abstract

Following events such as fatigue or stroke, individuals often move their trunks forward during reaching, leveraging a broader muscle group even when only arm movement would suffice. In previous work, we showed the existence of a ‘force reserve’ — a phenomenon where individuals, when challenged with a heavy weight, adjusted their motor coordination to preserve approximately 40% of their shoulder’s force. Here, we investigated if such reserve can predict hip, shoulder, and elbow movements and torques resulting from an induced shoulder strength deficit. We engaged 20 healthy participants in a reaching task with incrementally heavier dumbbells, analyzing arm and trunk movements via motion capture and joint torques through inverse dynamics. We simulated these movements using an optimal control model of a 3-degree-of-freedom upper body, contrasting three cost functions: traditional sum of squared torques, a force reserve function incorporating a nonlinear penalty, and a normalized torque function. Our results demonstrate a clear increase in trunk movement correlated with heavier dumbbell weights, with participants employing compensatory movements to maintain a shoulder force reserve of approximately 40% of maximum torque. Simulations showed that while traditional and reserve functions accurately predicted trunk compensation, only the reserve function effectively predicted joint torques under heavier weights. These findings suggest that compensatory movements are strategically employed to minimize shoulder effort and distribute load across multiple joints in response to weakness. We discuss the implications of the force reserve cost function in the context of optimal control of human movements and its relevance for understanding of compensatory movements post-stroke.

**NEW & NOTEWORTHY:** Our study reveals key findings on compensatory movements during upper limb reaching tasks under shoulder strength deficits, as observed post-stroke. Using heavy dumbbells with healthy volunteers, we demonstrate how forward trunk displacement conserves around 40% of shoulder strength reserve during reaching. We show that an optimal controller employing a cost function combining squared motor torque and a nonlinear penalty for excessive muscle activation outperforms traditional controllers in predicting torques and compensatory movements in these scenarios.

## INTRODUCTION

The redundancy of the human musculoskeletal system offers a multitude of potential solutions to even basic motor tasks, such as reaching a target with the hand (1). According to optimal control theory, the central nervous system chooses strategies that meet task requirements while minimizing effort (2). However, the precise definition of ‘effort’ in motor control is still a topic of ongoing debate (3–7). Within the optimal control framework, ‘effort’ is often quantified using cost functions, which serve as the optimization objectives guiding the selection of these motor strategies. In the present study, we propose and assess a new cost function rooted in foundational physiological principles, the reserve cost function, to refine the precision of motor command predictions, particularly in scenarios of reduced strength.

Fundamental neurophysiological principles are consistent with the “traditional” cost function of minimization of effort across joints. Henneman’s size principle suggests that motor units are activated in a sequence according to their force output, prioritizing smaller, more efficient, and less fatigable units (8). In addition, the temporal recruitment principle establishes that increasing force is associated with rising motor unit firing rates, thereby augmenting energy expenditures (9, 10). These principles highlight the progressive resource demands as the normalized muscle activation escalates. By penalizing higher muscle activation, the Central Nervous System (CNS) promotes workload distribution amongst effectors, leading to the formation of muscle synergies. This concept can be illustrated by the coordinated action of shoulder flexion and elbow extension during target-reaching tasks. Thus, instead of maximizing effort in one primary muscle, the CNS engages multiple muscles, allowing for shared effort and thereby conserving energy, reflecting a consistent and efficient CNS approach (11). In addition, muscles with similar activation percentages but different volumes may exhibit varying energy demands. A muscle of a larger volume, such as the gluteus, inherently consumes more energy than a smaller counterpart, like the anterior deltoid. In light of this, recent studies advocate for adjusting muscle activation based on muscle force or volume to enhance the prediction of motor commands (12). This logic aligns computationally with strategies that prioritize the minimization of the sum of the squared joint torques. Such strategies have been recognized for their association with energy consumption and robustness in predicting motor commands across diverse tasks (3, 11, 13, 14).

However, recent work suggests a need to modify this traditional cost function, at least when high torque levels are needed to accomplish the task. Individuals exhibit adaptive movement behaviors in situations where muscle strength or joint force is compromised. In seated reaching tasks, compensatory strategies like increased trunk recruitment appear in conditions such as post-stroke (15, 16), arm fatigue (17), or augmented arm weight (18). Notably, while individuals possess the capability to reduce trunk compensation, they opt against it, even when facing the increased mechanical demands of the compensatory strategy. This heightened trunk compensation, often at the expense of reduced shoulder-elbow utilization, is termed Proximal Arm Non-Use (PANU) in the stroke rehabilitation field (19). This suggests that the mechanical costs introduced by these adjustments can sometimes outweigh the strict biomechanical requirements for efficient movement (18, 19). Traditional explanations, which attribute excessive trunk compensation to post-stroke motor deficits (20, 21) and altered self-perceptions of capability (22, 23), might be insufficient. The manifestation of these compensations across both post- stroke and healthy populations suggests a shared underlying mechanism. Furthermore, whereas the traditional cost function predicts maximal muscle activation under high force demands, complete simultaneous activation of all muscle fibers is generally unattainable under normal physiological conditions (24). In addition, high-intensity tasks trigger fatigue beyond energy costs, implicating central mechanisms and metabolite accumulation (24–26). This introduces an interesting proposition: Could the CNS have a fail-safe mechanism penalizing extreme muscle activations to prevent early fatigue? In prior work (18), we showed the existence of a ‘force reserve’ — a phenomenon where individuals, when challenged with a heavy weight, adjusted their motor coordination during reaching movements to ensure that they preserved a reserve equivalent to approximately 40% of their shoulder’s maximum voluntary torque (MVT). In this study, we aim to evaluate the effectiveness of a new cost function based on the concept of a force reserve. Its performance is assessed against conventional predictive models for motor commands.

Here, we use a combination of behavioral and simulation methods to study reaching movements with induced shoulder strength deficits. We instructed young healthy participants to reach for a target positioned at arm’s length in front of them while holding dumbbells of increasing large weights. Arm and trunk movements were recorded using motion capture; joint torques were estimated via inverse dynamics methods. Simulations results of optimal controllers of a 3DOF trunk and arm planar model were contrasted with experimental results. We tested two main hypotheses: First, as we increase the weights during reaching tasks, there will be a proportional increase in trunk involvement to ensure that shoulder torque remains below 60% of its maximum capacity, thus conserving a force reserve close to 40%. Second, an optimal controller based on the new cost function of squared motor torque augmented with a nonlinear penalty that limits maximum contraction (reaching an asymptote at 100% MVT) will surpass controllers with the traditional cost function as well as that with another cost function with normalized torques in predicting compensatory movements and shoulder torques for large weights.

## MATERIALS AND METHODS

### Experimental Section

We extended our prior seated reaching task experiment involving healthy individuals (18) by introducing an expanded set of dumbbell weights. These weights comprised a control condition (no dumbbell), 0 kg (light foam dumbbell), 3 kg, 6 kg, 9 kg, 12 kg, 15 kg, and 18 kg. Due to safety concerns and personal comfort, results from three participants who could not, or chose not to, perform tasks with 15 kg and 18 kg weights were omitted.

### Participants

Twenty-one healthy subjects were initially screened for this study. One individual was excluded during the screening process, resulting in twenty participants (right-handed, 12 males, age 22 ± 8 years, height 1.73 ± 0.11 m, weight 66.8 ± 10.41 kg). We excluded individuals experiencing shoulder pain or any condition that might affect the reaching task. All participants provided their written informed consent before the study began. Our research complied with the 1964 Declaration of Helsinki and secured approval from the EuroMov Institutional Review Board at the University of Montpellier (IRB-EM 2106E).

### Procedure

Participants were instructed to touch a target, represented by a table tennis ball, using the side of their thumbnail. This ball was positioned at the end of an 80 cm high metal rod in front of the participant (19) (Figure 1). We adjusted the target’s distance to align with the participant’s fully outstretched arm, measured from the medial axilla to the distal wrist crease. During these measurements, participants were seated with their trunks resting against the backrest of the chair to prevent any trunk rotation.

**Figure 1.**
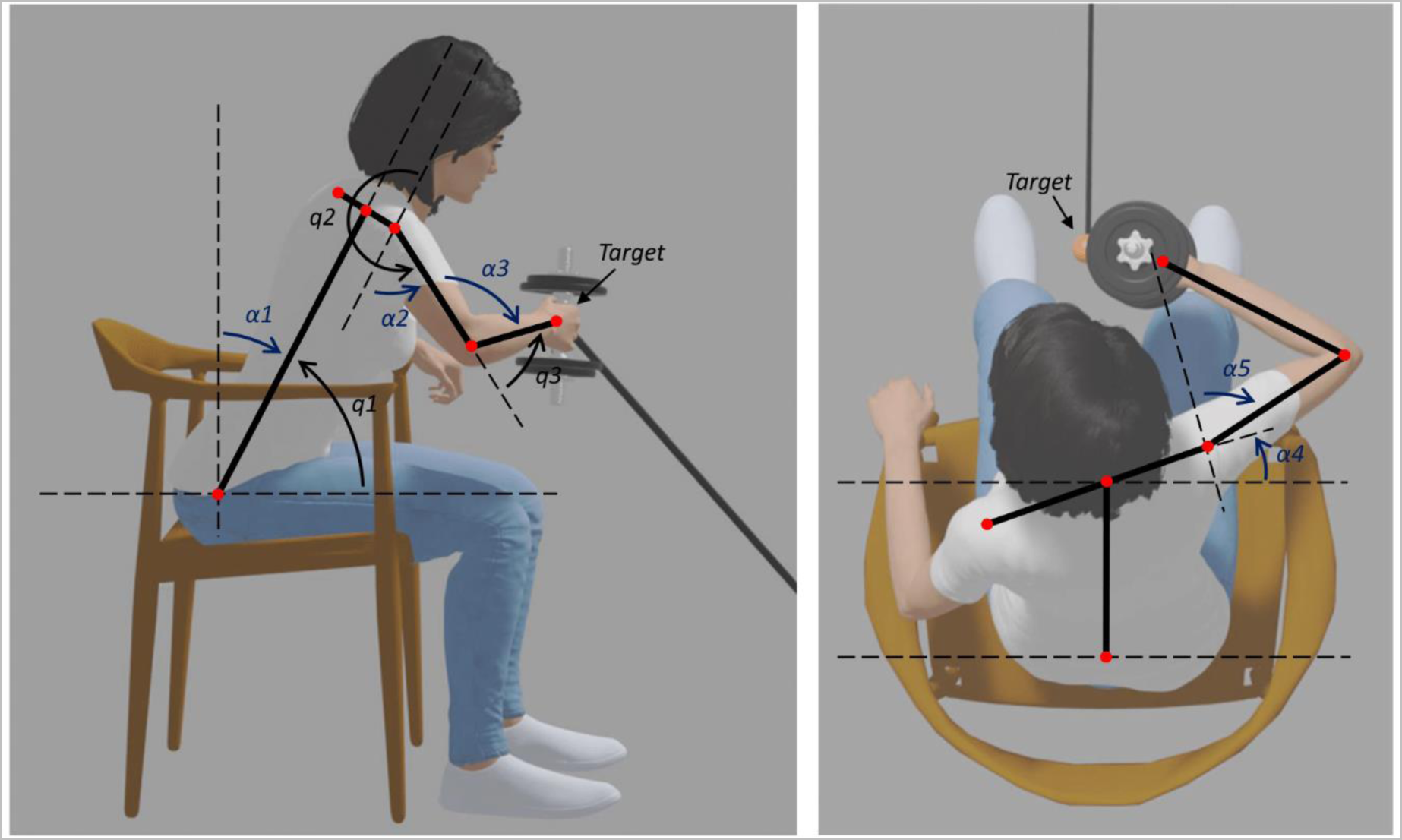
Experimental Set-Up and Angle Convention Overview. The left panel illustrates the sagittal plane, while the right panel presents the transverse plane. Participants, while seated, aimed to reach a target using their hands, holding a dumbbell. We assessed the body’s final posture (hand proximate to the target) via motion capture. Angles in black (q1, q2, q3) signify trigonometric angles for simulation calculations, whereas angles in blue (α1, α2, α3, α4, α5) denote the anatomical angles used in this manuscript for clarity. Specifically, for the simulations, q1, q2, and q3 represent hip, shoulder, and elbow flexions, respectively. For the anatomical angles, α1, α2, α3, α4, and α5 represent trunk flexion, shoulder flexion, elbow extension, trunk rotation, and shoulder abduction, respectively. Please note that angles α4 and α5 were exclusively descriptive and not incorporated in the minimal 2D-3DOF simulation.

Participants began seated with their feet flat on the ground and their backs against the chair’s backrest. Their forearms were comfortably placed on the armrests. They were instructed to naturally reach the target, pause for a second, and then revert to the starting position. Every participant performed three trials for each weight condition with their right hand, and the trial with the median compensation was retained for analysis. We chose the trial with the median trunk angle to ensure biomechanical validity, as mean values for posture analysis could result in non-viable biomechanical positions due to the averaging of angles (trunk, shoulder, elbow). This approach ensures our analysis reflects a physically attainable posture. We randomized the task sequence across all dumbbell weights such that all sequences of weights were different across participants. Breaks were interspersed between conditions to minimize fatigue.

To assess the shoulder’s maximum voluntary torque (MVT), participants, while seated, extended one arm in front of them, holding a 2 kg dumbbell. They were instructed to lift the dumbbell against gravity with maximum effort for 5 seconds. The dumbbell was connected to a ground-based force sensor via a static rope. Continuous backrest contact was maintained to ensure consistent posture and use of the opposing arm was restricted. Participants received consistent verbal encouragement from the experimenter. The maximum value from three separate trials, interspersed by one-minute rest intervals, was recorded. The highest recorded force was then employed to estimate the maximum shoulder torque, considering the participant’s arm posture and the torque induced by arm weight.

### Experimental Setup

Movements were recorded at 100 Hz using eight infrared optical cameras from the Vicon motion capture system (Vantage V5, 8.5 mm lens, Oxford Metrics, UK). Data from each marker were acquired using VICON Nexus 2 software. Markers were strategically placed on the target, manubrium, xiphoid process, both sides of the first metacarpal, styloid processes of the radius and ulna, medial and lateral epicondyles of the humerus, acromion process, and the iliospinale. For enhanced accuracy in data analysis, the positioning of the markers at the iliospinale was adjusted to correspond more closely with the anatomical hip joint center.

### Data Analysis

Data analysis was performed using Scilab 6.0.2.

#### Motion Capture

All position time series were filtered using a 5 Hz low-pass second-order Butterworth double-pass filter. Movement onset (t_0_) was determined when the 3D hand velocity within the workspace became positive and remained so until its maximum. Movement ending (*t*_final_) was determined once the Euclidean target distance reached its minimum, following Bakhti *et al*. previous work (19). We evaluated terminal posture at *t*_final_, with angle computations adhering to the conventions outlined in Figure 1.

#### Normalized Dumbbell Weight

To account for the variance in effort exerted by different participants when lifting a given dumbbell weight, we normalized each dumbbell weight. This was achieved by calculating the shoulder torque necessary for a participant to maintain the dumbbell near the target with their arm fully extended at the shoulder and elbow (a hypothetical value). This value was then divided by the participant’s maximum voluntary torque (MVT) at the shoulder. We refer to this metric as “Normalized Dumbbell Weight”, expressed as a percentage of the shoulder’s maximum voluntary torque (% shMVT).

### Simulation

#### Overview

We simulated the experimental task using an optimal control framework in which the body posture at target acquisition minimizes a cost function that encapsulates fundamental principles of muscle activation and torque production. We tested the ability of different cost functions derived from different sets of fundamental principles to predict joint positions and torques as the load at the hand increases (see *Cost functions* subsection).

We aimed to minimize the complexity of the model while capturing relevant aspects of the seated reaching task, especially trunk compensation in trials with a large load. Therefore, we modeled only the upper body, as the lower body is not primarily involved in the task. The model represents the upper body as a serial manipulator restricted to the mid-sagittal plane, thus capturing the task from a lateral perspective (Figure 1). The manipulator consists of 3 segments: trunk, upper arm, and forearm. This 3- DOF representation in the sagittal plane encapsulates the majority of movement organization inherent to the task. The trunk segment is attached to the ground frame by a revolute joint representing the hip joint (q_1_), allowing the trunk to lean forward or backward as in a seating posture. The upper arm segment is attached to the trunk segment by a revolute joint representing the shoulder joint (q_2_). Finally, the forearm segment is attached to the upper arm segment by a revolute joint representing the elbow joint (q_3_). The free end of the forearm segment is the end-point of the manipulator and represents the position of the hand/load.

As in the experimental task, loads of different weights are applied at the end-point of the manipulator. The objective of the task is to place the end-point of the manipulator at the target position for each load. The target position is a point in the mid-sagittal plane. Because the model of the upper body consists of 3 bodies moving in a two-dimensional space, there is, in general, an infinite number of configurations of the manipulator’s joints (postures) that satisfy the target position. Therefore, the CNS must select a single posture that is optimal according to the principles captured by the selected cost function.

### Cost Functions

We tested three different cost functions that encapsulate different principles of muscle activation and force production.

#### Traditional Cost Functions

The traditional cost function *C*_tra_ seeks to minimize effort, which is given by the sum of squared joint torques for static postural tasks (3, 11, 13, 14).

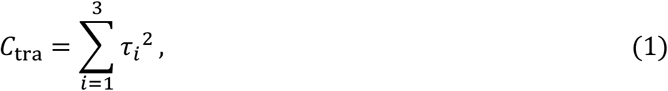

where *τ_i_* denotes the magnitude of the torque at the *i*-th joint, with *i* = 1, 2, 3 representing the hips, shoulder and elbow, respectively.

#### Augmented Torque: Incorporating Reserve Penalty

Recognizing that traditional cost functions do not account for physiological limits on muscle activation, we added a penalty informed by the force reserve to the traditional cost function (eq. 1). The force reserve *τ*_res,*i*_ indicates the capacity of joint *i* to generate additional torque given the current torque *τ_i_*:

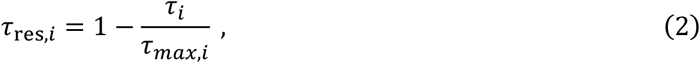

where *τ*_max,*i*_ is the maximal torque at joint *i*. The core idea of the reserve penalty is that it increases the cost as the force reserve diminishes, with an infinite cost aligning with tetanic contraction. Thus, we defined the reserve penalty *p_i_* as

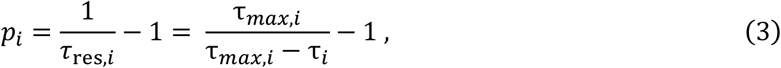

where we have used eq. 2. Notice that we have constructed the reserve penalty *p_i_* such that it approaches infinity as *τ_i_* approaches *τ*_max,*I*_, and *p_i_* = 0 when *τ_i_* = 0. However, while it is speculated that larger muscles might accumulate more fatigue than smaller muscles at similar activations, it is established that maximal force at each joint is related to the underlying muscle volume (27–29). Hence, we incorporated a scaling factor:

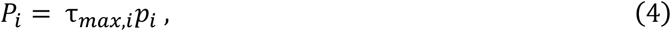

where, *P_i_* is the scaled reserve penalty at joint *i*. Incorporating the scaled penalty to the traditional cost function yields the augmented torque cost function *C*_res_:

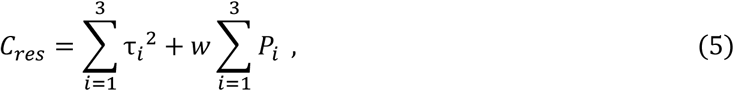

where *w* is a constant that weighs the contribution of the reserve penalty cost with respect to the traditional cost. Therefore, this function encompasses the duality of energy expenditure and fatigue minimization. Initially, to test our predictions, the performance of the reserve function was compared to other cost functions with *w* =1. Subsequently, simulations were conducted to find the optimal weight *w* of the force reserve that minimizes torque errors. The optimal value of *w* was determined using the Leave-One-Out Cross-Validation (LOOCV) method across all simulated subjects (see Fig. 4B). Specifically, for each iteration of LOOCV, one simulated subject was excluded from the dataset, and the model was trained on the remaining subjects. This process was repeated for every subject, and the optimal model weight was identified based on the lowest mean relative root mean square error (RMSE) across the training set. The final model weight was determined by averaging the best weights obtained from each LOOCV iteration.

#### Normalized Torque

We also test a recently proposed cost function, according to which humans minimize the effort exerted at each joint with respect the maximal effort value (12). Specifically, the torques produced at each joint arise from the squared activations of muscles normalized to their maximum potential activation (12). For a static postural task, the normalized torque cost function *C*_nor_ is:

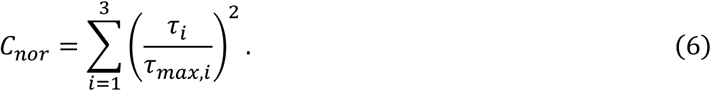

### Problem Definition

The optimization problem for the final posture in the seated reaching task can be expressed as

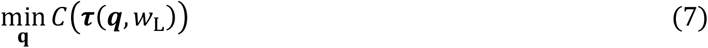

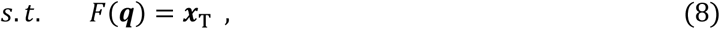

where *C* is a cost function of the joint torques ***τ***, which are in turn a function of the joint configuration ***q*** and the weight of the load *w*_L_. The form of the different cost functions that we used is described in detail in the *Cost functions* subsection. ***τ*** and ***q*** are vectors containing the torque and angle of each joint, respectively ([*τ*_1_ *τ*_2_ *τ*_3_]^T^ and ([*q*_1_ *q*_2_ *q*_3_]^T^). The solution of the optimization problem must satisfy the equality constraint in eq. 8, where *F* is the forward kinematics of the end-point of the manipulator and ***x***_T_ is the position of the target in cartesian coordinates. Thus, this constraint defines the target position of the end- point of the manipulator in the workspace.

To evaluate cost function *C* at any posture ***q*** it is necessary to compute the joint torques ***τ*** at posture ***q***. We used the principle of virtual work to compute ***τ***. First, we compute the joint torques ***τ***_L_ that counteract the external load at ***q***,

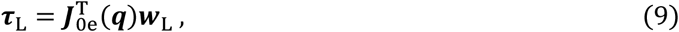

where ***J***_0e_ is the Jacobian matrix relating joint velocities to end-point velocity with respect to the ground frame in the manipulator (see *Appendix*), and ***w***_L_ is a force vector opposite in direction to the vector associated to the weight of the load. Therefore, ***w***_L_ = [0 *w*_L_]^T^, where *w*_L_ is the weight of the load. Notice that ***J***_0e_ is a function of the joint configuration ***q***. Next, we also compute the joint torques ***τ***_w*i*_ that counteract the weight of each segment *i* in the manipulator. These are obtained in a similar way:

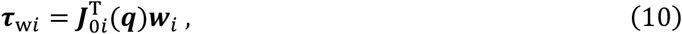

where ***J***_0*i*_ is the Jacobian matrix relating joint velocities to the velocity of the center of mass of segment *i* with respect to the ground frame (see *Appendix*), and ***w****_i_* is a vector opposite in direction to the vector associated with the weight of segment *i*. Here, *i* = 1, 2, 3 indicates the trunk, the upper arm, and the forearm, respectively. Therefore, the net torque at each joint is

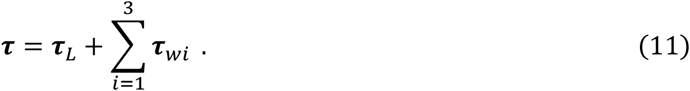

Notice that eq. 11 and eq. supp. 1 compute the torques at each joint using the same principle. However, in eq. supp. 1 the result is a 3D torque vector at a single joint, whereas in eq. 11 the result is a vector of 1D torques at all joints. This is because our simulations are based on a planar model of the upper body. Thus, in the simulations, torques are always orthogonal to the plane of the model.

### Optimization Procedure

We used a simple procedure that, for a given external load *w*_L_, allowed us to express the value of cost function *C* as a function of only *q*_1_. Namely, we set *q*_1_, the hip angle, as an independent variable that we swept to find the optimal posture ***q***. For a given angular position *q*_1_, the position of the trunk is fixed, and the position of the shoulder joint in the plane is immediately determined. This way, the upper arm and forearm segments can be seen as a reduced manipulator with two links, with the shoulder joint as the attachment to the ground frame. Using the inverse kinematics of this two-link planar manipulator, we can determine the angular position of the shoulder *q*_2_ and the elbow *q*_3_ (see *Appendix*). In general, for a given *q*_1_, there are two possible configurations for *q*_2_ and *q*_3_. However, we only consider the configuration where the upper arm points downwards. Therefore, for all swept values of *q*_1,_ we found a corresponding value of *q*_2_ and *q*_3_. This allowed us to compute the net torque ***τ*** at each joint configuration ***q*** for load *w*_L_ (eq. 11) and, thus, the value of cost function *C* (eq. 7, 1, 5, 6). For our analysis, we defined a range for *q*_1_ from 0 to π, using increments of π/1000. Within this range, finding the joint configuration ***q*** that minimizes *C* is straightforward. Thus, our model can predict the posture ***q*** of the upper body given an external load *w*_L_.

We created simulated subjects based on the experimental subjects tested in the study. Namely, for each simulated subject, the parameters in the model were taken from measurements of a real subject. These parameters are body segment lengths *l_i_*, target position **x**_T_, and maximal shoulder torque *τ*_max,2_ (see *Procedure* subsection in *Experimental Section*). Additionally, segment weights **w***_i_* and maximal hip and elbow torques (*τ*_max,1_ and *τ*_max,3_) were estimated from each participant’s height and weight. Torque values were adjusted using age- and gender-specific muscle strength ratios from Danneskiold-Samsøe *et al*. (30), which assessed isometric and isokinetic strength across major joints in a healthy population. The simulations were performed using R 4.2.2 and the pracma package (31).

### Model Validation and Statistical Analysis

The model’s fidelity was assessed by contrasting its predictions with experimental outcomes. The accuracy of the model was measured using the root mean square error (RMSE) for the discrepancies in joint angles and torques between the model’s predictions and the experimental data for each participant across different cost functions. For a concise presentation, a total RMSE was calculated for each participant by averaging the errors across the hip, shoulder, and elbow joints. Data outcomes are presented as mean ± standard deviation (SD).

Considering that the proposed augmented torque cost function incorporates an additional free parameter compared to the traditional and normalized torque cost functions, the Bayesian Information Criterion (BIC) was computed for all models. This metric evaluated the trade-off between model fit and parsimony.

To determine the optimal *w* in eq. 5 (the free parameter in the augmented torque cost function), we employed a two-stage approach: Initially, we explored a logarithmic scale spanning powers of 10 from 0 to 4 with 200 points. Subsequent refinement was conducted in a linear scale, specifically ranging from 10.6 to 10.8 in increments of 0.1. The value of *w* that produced the minimum RMSE, validated using the LOOCV method, was selected. A detailed sensitivity analysis of this parameter search is provided (Fig. 4B).

Statistical computations were performed using R version 4.2.2. Data visualizations were generated using the ggplot2 package (v3.4.1) (RRID:SCR_014601) (32).

## RESULTS

### Theoretical Predictions: Comparative Analysis of Postural and Torque Responses Across Models

Our force reserve model offers new theoretical predictions, which we compared with those from traditional and normalized models. We focused this comparison on postural adjustments and torque distribution, influenced by varying dumbbell weights during a simulated reaching task.

#### Posture Analysis: Model-Specific Trunk Flexion Responses to Varying Dumbbell Weights

Our analysis revealed that all three models—the reserve, traditional, and normalized—predict an increase in trunk involvement as the dumbbell weight increases. Significant differences among the models were most apparent at heavier weights, as shown in Figure 2. At lighter weights, the traditional and reserve models indicated minimal trunk engagement, primarily managed through shoulder and elbow movements. As the weights increased, each model exhibited distinct trunk flexion responses. The reserve model demonstrated a more pronounced increase in trunk flexion and a corresponding reduction in shoulder and elbow extension at the heaviest weights. The normalized model consistently predicted stronger trunk engagement across all weights, markedly diverging from the other models by overestimating trunk movement, especially at lighter weights.

**Figure 2.**
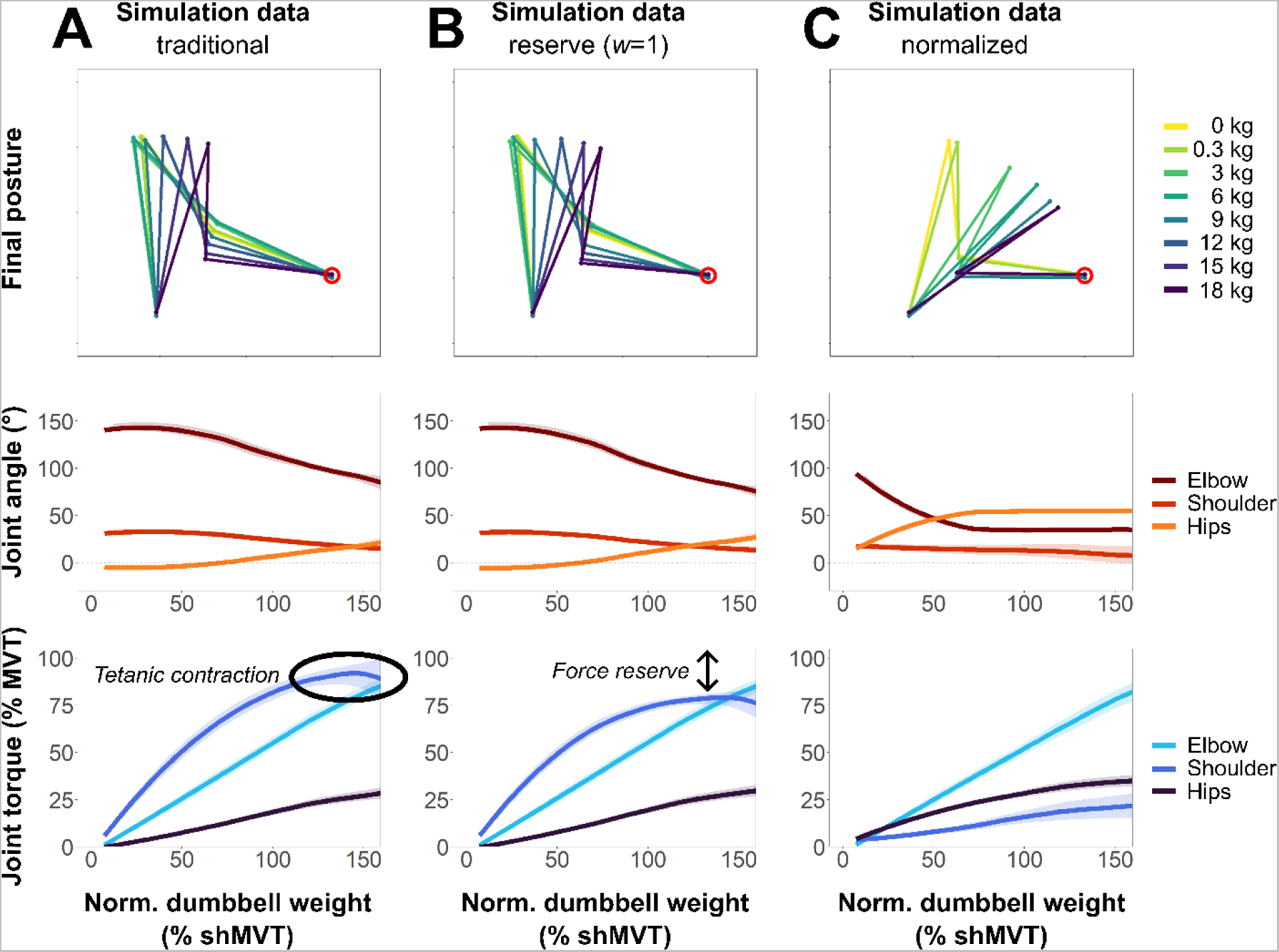
Comparative Analysis of Model Simulations Across Various Dumbbell Weights. This figure organizes the data into columns representing different models: Column A shows simulations using the traditional function, Column B those using the reserve function, and Column C those employing the normalized torque function. The first row illustrates 2D projections of a representative participant’s final posture, segmented into trunk, upper arm, and forearm plus hand, with the target position marked by a red circle. Notably, as the arm weight increases, all models demonstrate increased trunk movement. However, significant differences in arm recruitment and its consequent impact on shoulder flexion and elbow extension are evident between the models. The second row displays flexion angles for the elbow (in dark red), shoulder (in light red), and hips (in orange), plotted against the normalized dumbbell weight, expressed as a percentage of the participants’ shoulder maximum voluntary torque (% shMVT). Thick lines represent the mean across simulated subjects, while shaded areas indicate the 95% confidence interval. As the dumbbell weight varies, shoulder and elbow angles adapt, leading to augmented trunk involvement. Although the models share qualitative similarities, they differ quantitatively. The third row presents torque values for the elbow (in light blue), shoulder (in blue), and hips (in dark blue) in relation to dumbbell weight. The traditional model often exhibits tetanic contraction of the shoulder muscle, with shoulder torque frequently reaching its maximum. In contrast, the reserve model shows a threshold of shoulder torque below 80% MVC, consistent with the force reserve hypothesis. Meanwhile, the normalized torque model demonstrates significant trunk recruitment, resulting in correspondingly reduced torque at the shoulder joint.

#### Torque Analysis: Variations in Shoulder Torque Thresholds in Traditional versus Reserve Models

Our torque distribution analysis across the elbow, shoulder, and hip joints revealed that both the traditional and reserve models initially show a near-linear increase in shoulder torque up to about 60% of maximal voluntary torque (shMVT), as detailed in Figure 2. At lighter loads, torque values are minimal, increasing significantly as the dumbbell weights rise to 40-60% shMVT. Beyond this point, the models diverge notably. The traditional model continues to show an increase in shoulder torque, peaking at 90% MVT for the heaviest weights, with 8 out of the 20 simulated participants surpassing 99% MVT, indicating tetanic muscle contractions. In contrast, the reserve model displays a plateau in shoulder torque below 80% MVT, demonstrating the impact of the reserve force penalty. The normalized model significantly differs, showing higher trunk torque from lighter weights and a much slower increase in shoulder torque, peaking at only 21% MVT for the heaviest weights, as seen in Figure 2.

### Experimental Results: Observed Motor Adaptations to Varied Load Conditions in Healthy Participants

Our experimental investigations revealed detailed insights into the postural adjustments and torque distributions observed in participants performing reaching tasks with varying dumbbell weights. This analysis serves as a foundation for understanding actual motor behavior in the context of our theoretical models.

#### Posture Analysis: Graduated Shifts in Participant Postures with Increasing Weights

We observed how participants adapted their postures in response to incremental dumbbell weights, reflecting physical adjustments to increasing loads. These adaptations involved changes in shoulder flexion, elbow extension, and hip positioning, as depicted in Figure 3A. Initially, for lighter weights (≤ 10% shMVT), the posture primarily involved shoulder flexion and elbow extension with slight hip extension to counterbalance the trunk torque. As weights increased to 40-60% shMVT, participants began altering their movement strategies, showing a minor transition in all joint angles. With even heavier weights (90 to 110% shMVT), trunk involvement became more significant, and joint angles in shoulders and elbows notably decreased. At the highest weight range (140 to 160% shMVT), measurements indicate substantial hip flexion with marked reductions in shoulder flexion and elbow extension, aligning with the physical demands of handling heavier loads.

**Figure 3.**
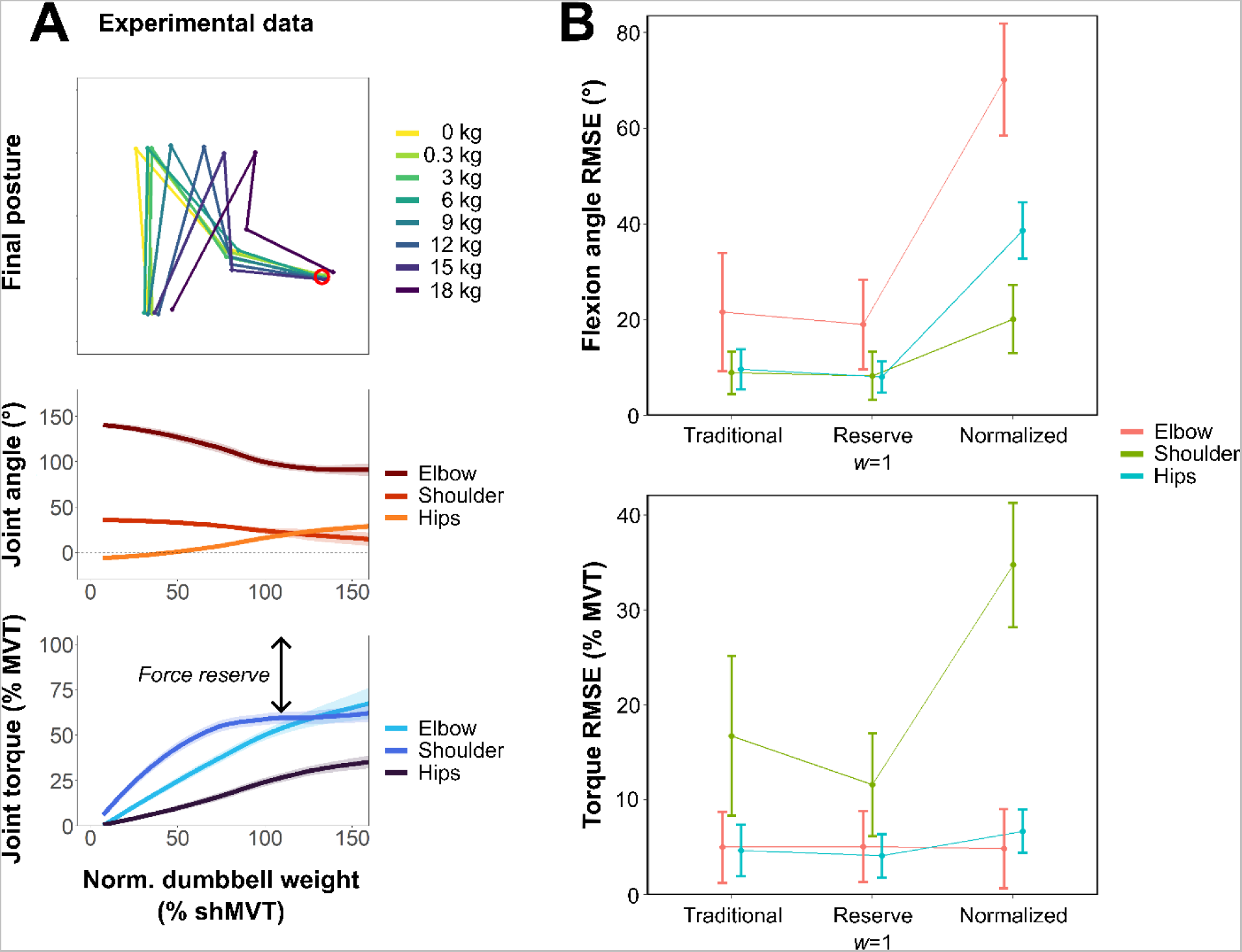
Comparative Analysis of Experimental Data and Model Simulations. Panel A presents an analysis of experimental data across various dumbbell weights. The first row depicts 2D projections of a representative participant’s final posture, segmented into trunk, upper arm, and forearm+hand across diverse dumbbell weights, with the target position denoted by a red circle. The upper arm length appears variable due to the 2D projection of our 3D model. As the arm weight increases, the dataset shows pronounced escalation in trunk movement. The second row depicts flexion angles for the elbow (dark red), shoulder (light red), and hips (orange) against the normalized dumbbell weight. These weights are expressed as a percentage of participants’ shoulder maximum voluntary torque (% shMVT). Thick lines represent the overall mean, with shaded areas indicating the 95% confidence interval. As dumbbell weight varies, shoulder and elbow angles adjust, necessitating augmented trunk involvement. The third row depicts torque values for the elbow (light blue), shoulder (blue), and hips (dark blue) in relation to dumbbell weight. Experimental findings reveal that shoulder and elbow torques intensify with the increasing dumbbell weight, plateauing around 60% MVT, while trunk torque increases more gradually. Panel B focuses on the comparison of RMSE values across prediction models for joint angles and torques. The dot represents the mean deviation from experimental data, and the bars represent its standard deviation between participants. Of all the models assessed, the reserve and traditional models demonstrated the lowest prediction errors for joint angles. However, the reserve model showed superior accuracy for shoulder torque predictions than the traditional and normalized torque models. Notably, the normalized torque model performed the least effectively in predicting angles and torques.

**Figure 4.**
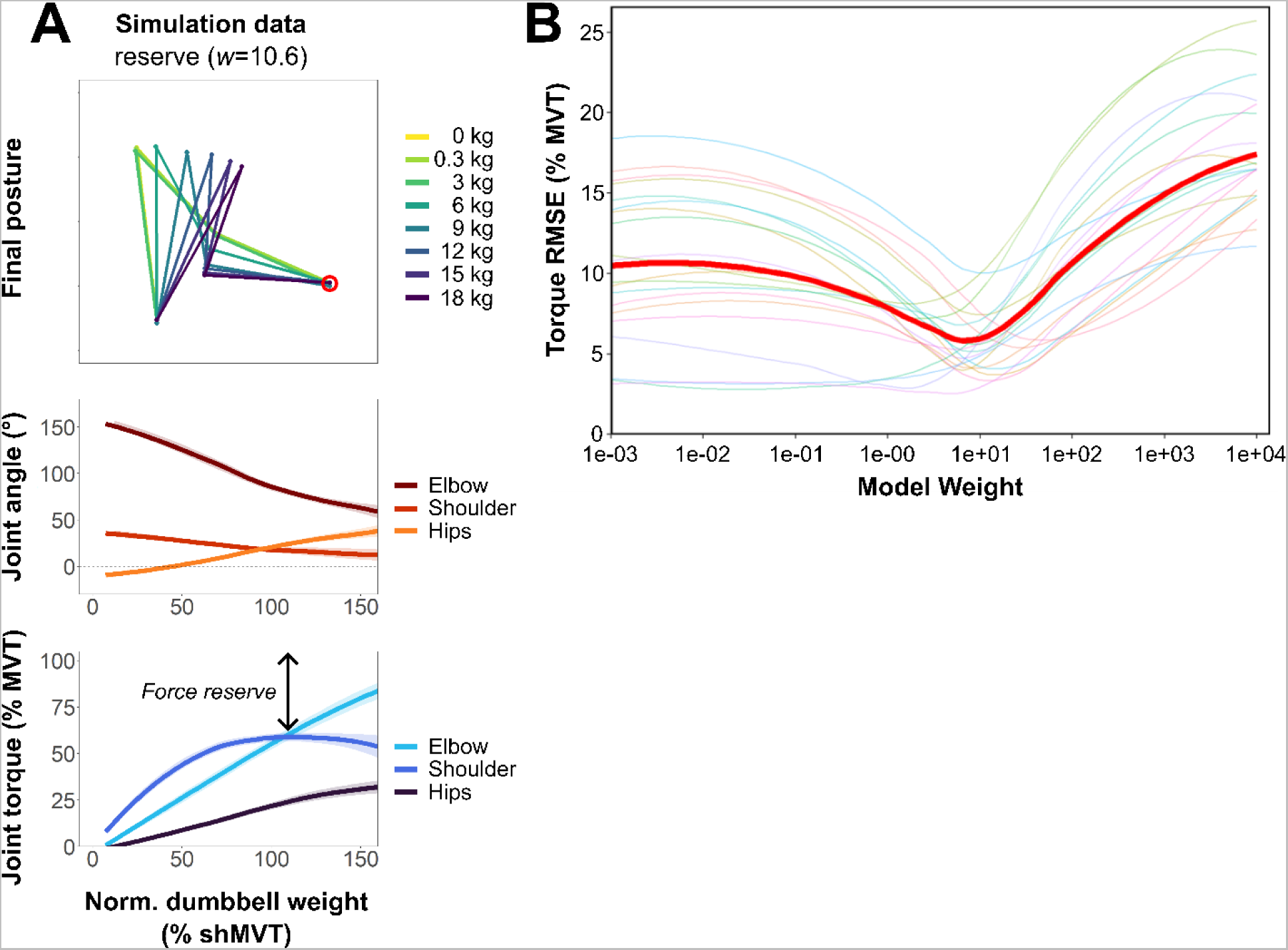
Analysis of Optimized Reserve Model Simulation. Panel A presents an analysis of simulation data of the optimized reserve model across various dumbbell weights. The first row depicts 2D projections of a representative participant’s final posture, segmented into trunk, upper arm, and forearm+hand, with the target position denoted by a red circle. As the arm weight increases, the dataset shows pronounced escalation in trunk movement. The second row depicts flexion angles for the elbow (dark red), shoulder (light red), and hips (orange) against the normalized dumbbell weight. These weights are expressed as a percentage of participants’ shoulder maximum voluntary torque (% shMVT). Thick lines represent the mean across simulated subjects, with shaded areas indicating the 95% confidence interval. Variations in dumbbell weight lead to adjustments in shoulder and elbow angles, resulting in increased trunk involvement. The third row depicts torque values for the elbow (light blue), shoulder (blue), and hips (dark blue) in relation to dumbbell weight. As in the experimental data, the simulation indicates that shoulder and elbow torques intensify with increasing dumbbell weight, plateauing around 60% MVT, while trunk torque rises more gradually. Panel B focuses on a sensitivity analysis of the parameter w. The log-scaled x-axis highlights variations in the w parameter, while the y-axis corresponds to overall mean RMSE values for torque estimation. Thin lines represent individual subjects; the thick red line shows the smoothed average. The graph emphasizes the significance of the optimal w value (10.6) in enhancing model accuracy. When the w value approaches 0, the reserve penalty becomes negligible, mirroring the traditional function. Such findings emphasize the clear superiority of the reserve function when incorporating an appropriate penalty term.

#### Torque Analysis: Experimental Observations of Shoulder Force Reserve at 60% Maximal Voluntary Torque

Our analysis of torque variations at the elbow, shoulder, and hip joints with increasing dumbbell weights demonstrated clear biomechanical adaptations as weights increased. Consistent with findings from Faity et al. (18), torque at these joints increased nearly linearly until reaching approximately 60% of maximal voluntary torque (MVT). For lighter weights, torque increments were modest but grew substantially as weights approached and surpassed 40% to 60% shMVT. The heaviest weight categories, particularly beyond 90% shMVT, saw the most significant increases. A distinct observation from our study was the early attainment of the 60% torque threshold by the shoulder joint. As this threshold was approached, there was a concomitant elevation in trunk involvement, suggesting a strategy to conserve shoulder force reserves as the dumbbell weight increased.

### Comparison with Simulation Results: Identifying the Most Accurate Model Predictions

We compared the simulated predictions from the traditional, reserve, and normalized models against experimental findings to identify the model that most accurately reflects real-world data. This evaluation is essential to validate the effectiveness of the reserve model in predicting joint angles and torques across different weights.

#### Posture Analysis: Superior Performance of Reserve and Traditional Functions Over Normalized Model

We evaluated the accuracy of each model in predicting participant postures with varying dumbbell weights. The traditional model yielded precise estimates of trunk movement for lighter weights but undershot it as the weight intensified (Figure 2A - Final posture and Joint angle). Contrarily, the reserve function deviated from the predictions of the traditional function, especially for the heaviest weights, showing a greater fidelity of hip flexion for medium to high dumbbell weights (Figure 2B - Final posture and Joint angle). Finally, the normalized torque function systematically overestimated trunk movement in all weight conditions, even without the dumbbell (Figure 2C – Final posture and Joint angle).

#### Torque Analysis: Reserve Function’s Superior Accuracy in Joint Torque Prediction

We evaluated the precision of each model in estimating the torque demands at the elbow, shoulder, and hips, particularly noting how well they represent the threshold of muscle activation and the concept of force reserve. The traditional cost function’s predictions displayed discrepancies with the experimental data for heavier dumbbell weights (Figure 2A – Joint torque), underestimating trunk movement and overestimating shoulder engagement. In contrast, the reserve function more accurately represented the threshold of shoulder muscle activation, attributable to its penalty term (Figure 2B – Joint torque). Nevertheless, it predicted a reserve threshold at around 80% MVT for the shoulder, compared to the experimental values of 60% MVT. The normalized torque function, as hypothesized, underestimated shoulder torque and overestimated hip torque (Figure 2C – joint torque).

#### Model Comparison: Reserve Torque Model as the Most Parsimonious and Accurate

The Bayesian Information Criterion was applied to the torque RMSE (% MVT) to determine the most parsimonious model among those evaluated. The resulting BIC scores were: −82.6 for the traditional model, −91.9 for the reserve torque model, and −56.2 for the normalized torque model. A lower BIC value indicates a better model fit when accounting for complexity. Of the three models, the reserve torque model exhibited the greatest balance between goodness-of-fit and parsimony, with a BIC difference of 9.3 compared to the traditional model. This difference provides strong evidence of its superiority, as a BIC difference ranging from 6 to 10 corresponds to a Bayes factor of 20 to 150 (33).

Additionally, we explored variations of the traditional cost function by increasing the exponent to 4 and 10, which yielded BIC scores of −78.7 and −77.6, respectively. These adjustments increased the penalty on larger torques but failed to consider the maximum torque capacity of each joint, leading to increased constraints on weaker joints like the shoulder. Consequently, these models showed less favorable fits compared to the reserve torque model, as indicated by their less negative BIC scores.

### Refining the Reserve Model: Fine-Tuning the Reserve Penalty Weight

We investigated whether fine-tuning the reserve penalty weight (*w*) in the reserve function could enhance simulation accuracy, particularly for shoulder torque, without compromising other aspects such as posture and torque at other joints. Our objective was to identify an optimal reserve penalty weight that minimizes torque errors.

#### Determining the Optimal Reserve Penalty: Sensitivity Analysis of Parameter w

We executed a sensitivity analysis focusing on the parameter *w* (eq. 5), which balances the effect of torque minimization and torque reserve. Using the LOOCV method with RMSE analysis, we found the optimal *w* value to be 10.6, with a 95% confidence interval of [10.4, 10.8]. As a *w* value close to 0 makes the reserve penalty negligible, reflecting the traditional function, our results corroborate the superiority of the reserve function when a penalty term is in play. The influence of changes of *w* on the model’s predictive accuracy is displayed in Figure 4B.

#### Enhanced Model Accuracy: Simulation Outcomes with Tuned Reserve Penalty

As shown in Figure 4A, fine-tuning the reserve penalty weight *w* further enhanced the accuracy in predicting posture and torques. This adjustment led to an accurate representation of reserve force, aligning the shoulder torque threshold with our hypothesis and experimental data at 60% MVT (Figure 4A).

Although the adjustment in reserve penalty weight *w* improved model precision, it also added complexity by introducing a new parameter. Consequently, we recalculated the Bayesian Information Criterion (BIC) to evaluate whether including parameter *w* compromises the model’s parsimony. The model with optimized *w* yielded a BIC of −103.2, compared to −91.9 without *w* optimization, suggesting that the optimized reserve torque model with *w*=10.6 is the most parsimonious among all tested models. This difference of 11.3 provides very strong evidence of its superiority, corresponding to a Bayes factor of over 150 (33).

### Simulation of Reduced Maximum Shoulder Torque

To further understand the potential compensatory mechanisms the central nervous system might adopt in the context of shoulder weakness, we simulated reaching movements across a spectrum of shoulder strength deficits, ranging from 0% to 99%. We designed these simulations with dual objectives: firstly, to evaluate the degree of trunk compensation relative to the varying levels of shoulder force deficit, and secondly, to investigate the occurrence of Proximal Arm Non Use (PANU), which is characterized by unnecessary trunk compensation during movement (19).

Each evaluated cost function indicates an increase in trunk involvement as shoulder strength wanes (Figure 5). However, the nuances of how each function responds to diminishing strength vary. A key distinction arises when juxtaposing these simulations with our previous set where dumbbell weight increased, but shoulder strength remained constant.

**Figure 5.**
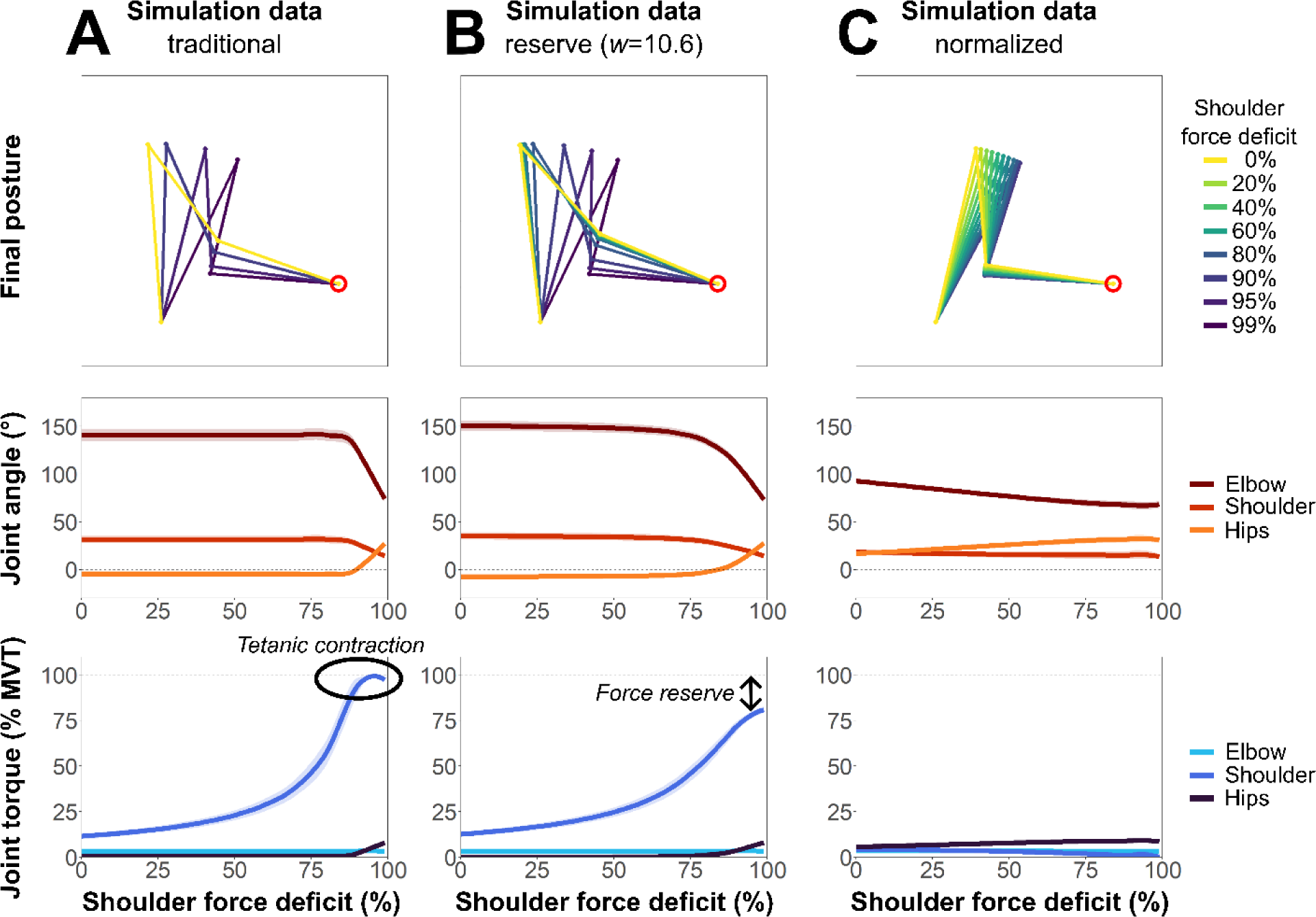
Comparative Analysis of Posture, Flexion Angles, and Torques in Response to Shoulder Weakness Across Different Cost Functions. The first row presents simulated postures of a representative participant, differentiated by degrees of shoulder weakness. The second row charts the elbow (dark red), shoulder (light red), and trunk (orange) flexion angles in relation to varying degrees of shoulder weakness. The third row illustrates the corresponding torques for the elbow (light blue), shoulder (blue), and trunk (dark blue) as they respond to shoulder weakness. For the second and third rows, thick lines show the overall average, whereas the shaded areas indicate the 95% confidence interval of the mean. The data is further categorized by columns, showcasing the outcomes for different functions. Column A elucidates the results derived from the traditional function. Column B focuses on outcomes from the reserve function (optimized with w=10.6). Column C shows the results using the normalized torque function. Notably, the traditional function maintains a consistent posture in relation to weight until the peak shoulder torque is attained, after which there is an obligatory shift in trunk movement. In contrast, the reserve function initiates a gradual increase in trunk movement, evident from around a 70% deficit. This movement intensifies as the degree of weakness escalates. The normalized torque function shows early and extensive trunk movement, which changes little as the strength deficit increases.

For the traditional function, the cost calculation remains unaffected by the maximum torque value, focusing solely on the joint torque limits. Consequently, alterations in maximum shoulder torque do not influence the movement strategy unless the torque breaches its joint-specific limit. As a result, the traditional function does not suggest a linear increase in trunk compensation. It only highlights the necessity for compensation when the force deficit is close to 90% (Figure 5A – joint torque). In cases of a 70% shoulder strength deficit, the traditional function does not show the anticipated increase in trunk compensation. This finding is in stark contrast with the higher trunk compensations typically associated with the PANU phenomenon at similar levels of deficit (34).

In contrast, both the reserve function and the normalized torque function integrate the maximum torque value in two ways: to set joint torque limits and to influence the movement’s cost. Thus, even if the maximum torque is not fully utilized, a reduction in its value can subtly shift the perceived cost of a movement. The reserve function reflects this nuance, suggesting a gradual increase in trunk compensation as shoulder strength drops (Figure 5B). It recognizes the possibility of PANU, given that the shoulder torque is not maximized. However, it slightly underpredicts compensation at a 70% deficit, hinting at other potential influencers of trunk compensation.

Lastly, the normalized torque function, while suggesting an increase in trunk compensation with waning shoulder strength (Figure 5C), still overpredicts compensation in standard force scenarios, suggesting it might do so in shoulder force deficit contexts as well.

## DISCUSSION

In this study, we show that increasing arm weight, resulting in a force deficit at the shoulder, significantly alters coordination in healthy participants. As this deficit intensifies, we noted an associated augmentation in trunk movement, evident through trunk flexion and rotation. Simultaneously, shoulder abduction increases while shoulder flexion and elbow extension decrease. These adjustments strategically position the arm’s center of gravity closer to the shoulder, effectively reducing the shoulder torque for a given dumbbell weight. These postural adjustments consistently maintain the shoulder torque within 60% of the maximum voluntary torque, preserving an approximate force reserve of 40% MVT. Our data confirm and extend the earlier insights on joint torque reserve presented in our prior research (18).

In accordance with our second hypothesis, our findings underscore the superior predictive efficacy of the reserve cost function, grounded in the foundational neurophysiological tenets of motor unit recruitment – even when accounting for added complexity. Furthermore, when simulating increasing shoulder weakness, the reserve cost function predicted both a progressive increase in trunk compensations and the phenomenon of PANU (due to never reaching maximum shoulder torque). These results highlight the role of the joint torque reserve in post-stroke compensation and nonuse.

### Trunk Compensation is an Optimal Solution to Alleviate Shoulder Weakness

#### In Healthy Participants

When arm weight is light in relation to available strength, reaching a target with only the arm facilitates movement by engaging minimal muscle mass, thus minimizing the mechanical cost. Under such conditions, healthy participants typically demonstrate reduced effort in their reach (Figure 3A). As the arm weight increases (e.g., with the addition of a dumbbell), limited trunk involvement continues to represent the most mechanically efficient approach (Figure 3B). Nonetheless, this situation necessitates heightened muscle activations. Importantly, muscle efficiency decreases as muscle activation intensifies, predominantly influenced by the spatial and temporal principles of motor unit recruitment (25, 35). Hence, as maximum muscle activation approaches, increasing effort is required for subsequent activation increments. Herein lies the value of recruiting additional degrees of freedom: it helps distribute the effort across multiple effectors (2, 11). The associated cost of engaging more substantial muscle groups is effectively offset by the efficiency gained from decreasing the normalized activation demand on each distinct muscle.

Furthermore, our results suggest that the mere minimization of energy requirements is insufficient to model human movement accurately. We show that the distribution of effort is wider, maintaining approximately 40% MVT as a force reserve across involved joints. This observation is consistent with the neurophysiological principles delineated in the introduction, proposing that the force reserve functions to prevent high-intensity muscle activations. Such activations can precipitate central and peripheral fatigue, which is influenced by spinal and supraspinal pathways, accumulation of metabolites, and diminished energy efficiency (24–26).

### When Facing Shoulder Weakness: Consequences for Stroke Patients

A stroke is a brain lesion that disrupts the normal signals sent to muscles, leading to diminished muscle activation. This disruption results in hemiplegia, a condition with a considerable drop in force, especially visible on one side of the body. While this impairment affects all muscles in the affected half of the body at varying degrees, its impact is most visible at the shoulder during upper limb tasks like reaching.

The guiding principle stays the same when shoulder strength diminishes, but arm weight remains unchanged (Figure 5). Efforts are channeled to minimize energy consumption while conserving some residual strength. This principle may explain the manifestation of trunk compensations and the tendency for PANU (Proximal Arm Non-Use, where heightened trunk compensation occurs at the expense of reduced shoulder-elbow utilization) as an energy conservation strategy. Such compensatory behaviors might provide insight into the observed trunk compensations in individuals with a previously fatigued arm (17) or stroke survivors (15, 16), where shoulder strength is compromised.

Shoulder weakness is the main contributor to upper limb functional deficits after stroke (36, 37). Such weakness has also been suspected as a potential cause of compensatory movement strategies (38). As illustrated in our findings (Figure 3, Figure 5), even when compensatory strategies are employed in a reaching task, the shoulder torque escalates more rapidly than that of the elbow and trunk. Since many daily activities revolve around hand movements for object manipulation, compromised shoulder strength can be a significant constraint (39). Furthermore, while compensating with the trunk is viable when the target is positioned below shoulder level, challenges arise when the target is higher. In such cases, patients might employ alternative compensatory mechanisms (like using their less affected arm) or struggle to execute the task successfully (40).

However, the reserve function fails to predict compensations substantial enough for detection when faced with a shoulder strength deficit of 70%, analogous to the average stroke patient (34). This suggests that the implications of shoulder weakness on effort may not be the only factor contributing to trunk compensation. Indeed, muscle strength is not the only determinant influencing stroke outcomes. Deficits in fine motor control, somatosensory functions, and occasionally the presence of spasticity or pain, impose additional constraints on the use of the paretic arm (41). These other deficits suggest that post-stroke compensations might not be exclusive to effort minimization. They could also serve other objectives, like enhancing the precision of arm movements via trunk control, among other reasons (16, 23). It is also plausible that a combination of these causes contributes to the emergence of compensations (42–44), in line with the composite cost function hypothesis (3).

### The Reserve of Joint Torque to Improve Effort Prediction

In the domain of biomechanical modeling and AI systems designed for human-like movement generation, the selection and refinement of cost functions play a pivotal role. The traditional cost function, predominantly focused on minimizing mechanical costs, has been instrumental in many optimal control studies and AI models (3, 11, 13, 14). Our results indicate that this function accurately predicts posture and motor commands at lower dumbbell weights.

However, the traditional function overlooks critical considerations. Specifically, it does not account for the heightened risk of fatigue—both central and peripheral—stemming from elevated muscle activation (24–26), nor for the inability to reach true maximal contraction (24). This oversight leads to an underestimation in the work distribution across involved effectors. Consequently, when the required normalized force is high—evident during tasks with heavier dumbbells or a shoulder strength deficit—the function underestimates trunk contribution and yields increased errors in motor command predictions. In contrast, our proposed penalty based on the force reserve principle balances energy expenditure and normalized muscle activation. It proficiently captures the aversion to high muscle activations, thereby ensuring the preservation of a joint torque reserve. Notably, when simulating a shoulder strength deficit, this function forecasts the proximal arm’s nonuse—a phenomenon aligning with observations in stroke patients (19).

Our findings are consistent with recent trends in gait research. Historically, while numerous cost functions have focused on arm control, both in biological and non-biological contexts (14, 45–49), gait control has primarily centered on minimizing metabolic energy (50–53). However, certain studies underscore a superior predictive accuracy when emphasizing reduced muscle activation (54, 55), thereby contesting the metabolic energy minimization paradigm. Contemporary evidence indicates a nuanced trade-off between metabolic energy and muscle activation reduction. This equilibrium, achieved via volume-weighted muscle activation (12), improves gait control predictions. Our results suggest that while energy minimization remains essential, it likely concurrently operates with another principle: moderating muscle activation to preserve a force reserve, which could serve as a protective buffer against unforeseen events and premature fatigue.

Extending these insights into AI-generated human-like movement, the significance of our reserve function becomes even more pronounced. In AI systems, where the goal is to replicate human biomechanics authentically, employing the traditional cost function might lead to strategies that, although efficient, are not reflective of genuine human movement patterns, particularly under high-load conditions. This mismatch can yield AI movements that are unrealistic or unsafe. Conversely, an AI trained with a reserve-based cost function learns to respect human movement’s inherent limitations and safety margins. This approach fosters AI-generated movements that are not just mechanically efficient but also safe, adaptable, and truly human-like. It is especially crucial in contexts where AI systems interact with humans, such as in assistive robotics or virtual human simulations (56).

### Limitations and Perspectives

#### Limitations of the Model

Our assumptions were tested on a simplified model (static, 2D, 3 DOF). This is a strength, as it demonstrates the robustness of the principles of joint torque reserve and work distribution across multiple effectors, and a limitation, as it moves away from the complexity of real-world scenarios. To test whether the principles described here are generalizable, it is necessary to test them in other tasks, using more ecological models and extending the cost function to dynamics.

Specifically, the optimized reserve function tends to overestimate hip flexion slightly under increased load conditions (Fig. 4 – Joint angle). Experimental data, on the other hand, reveal adaptive strategies employed by participants, such as enhanced leg recruitment, trunk rotation, and shoulder elevation (Supp. Data #1). These discrepancies may arise from the design constraints of our model, particularly the omission of leg movement, trunk rotation, and shoulder elevation. It should be noted that the observed overestimation may reflect real-world effort policies more accurately than the underestimation observed in the traditional function. This inference is corroborated by the superiority of joint torque predictions provided by the reserve cost function.

Furthermore, in our models, we observed a near-linear increase in elbow torque. This contrasts with the pronounced inflection present in the experimental data. This discrepancy may arise from constraints inherent to our model. Due to the imposed limitations—specifically, the inability to move the hips closer to the target or flex the elbow beyond 35°—the forearm was required to remain approximately parallel to the ground. This likely contributed to the observed quasi-linear trend in elbow torque.

Finally, despite the enhanced performance of our cost function when integrating the force reserve penalty, it is essential to clarify that this function is not designed to precisely replicate the actual cost function of the CNS. Instead, it aims to offer an approximation that errs the least and presents a more generalizable estimate.

#### Perspectives

While the joint torque reserve provides a compelling rationale for the compensations observed in stroke patients and healthy individuals with weighted or fatigued arms, it is essential to highlight that we have yet to experimentally test the reserve function directly in stroke participants—a definitive step required to validate this hypothesis. On a promising note, a clinical trial is currently underway (https://clinicaltrials.gov/ct2/show/NCT04747587). Preliminary findings from this trial suggest the following patterns in stroke patients: 1) trunk compensations and nonuse effectively reduce the required effort for movement compared to scenarios with minimal trunk engagement, as evidenced by the observed shoulder torque, muscle activation, and rate of perceived exhaustion, 2) reducing the arm’s weight leads to a reduction in these compensations, 3) a smaller target tends to elicit more pronounced compensations compared to a larger one (57). These initial results align with our hypotheses, further strengthening our position.

These clinical findings and the ‘Simulation of Reduced Maximum Shoulder Torque’ section illustrate that nonuse in stroke survivors can be partly explained by the force reserve principle, challenging the traditional view that nonuse is merely detrimental. This demonstrates that nonuse could be an efficient strategy, minimizing effort by avoiding muscle saturation, in contrast to ‘learned nonuse,’ where nonuse results from a misperception of capabilities. This suggests that tailoring rehabilitation strategies based on the force reserve phenomenon could significantly improve the decision-making process in assigning specific interventions for stroke patients. By utilizing the Proximal Arm Non-Use (PANU) test to assess compensatory trunk movements and the nonuse of the paretic arm, we could correlate these behaviors with functional outcomes and perceived effort. This assessment could allow clinicians to categorize patients into Optimal Compensators, who benefit from their current compensatory strategies, and Suboptimal Compensators, who do not benefit and are thus candidates for targeted interventions, such as Constraint-Induced Movement Therapy (CIMT) or trunk restraint, aimed at minimizing compensations. Furthermore, our approach emphasizes strengthening the antigravity muscles of the shoulder, major contributors to increased compensatory movements. Implementing specific resistance training with fewer repetitions and higher muscle activation should enhance shoulder strength (58–60) and might reduce reliance on compensatory movements. Integrating this strength training with therapies such as trunk restraint or CIMT could further reduce the dependency on compensatory strategies in daily activities. For patients with severe shoulder impairments who may not fully benefit from traditional strength training (61), deploying assistive technologies such as exoskeletons or passive lightening systems could be more appropriate (62). By focusing on supporting the shoulder joint during various tasks, these technologies could reduce the load on weakened muscles, thus facilitating arm use. These preliminary insights are part of an ongoing exploration, with more detailed findings and clinical implications to be discussed in an upcoming article focused on clinical results.

### Conclusion

Our research underscores the pivotal role of joint torque reserve in guiding compensatory strategies among healthy participants, with potential repercussions for stroke patients, especially as shoulder force diminishes. We introduced a novel cost function, which adeptly integrates the principles of force reserve, and demonstrated its superior predictive efficacy over traditional models. This reflects the balanced objective of minimizing energy while safeguarding against excessive muscle activation. Although our findings are rooted in a simplified 2D model, they shed light on the fundamental neurophysiological principles governing motor unit recruitment, offering insights into arm control and possible post-stroke compensations. Future research should delve into more complex and ecological models to validate and extend these findings.

## APPENDIX

## Jacobian Matrices

The Jacobian matrices relating joint velocities to velocities at the end-point and the center of mass of each link with respect to the ground frame are

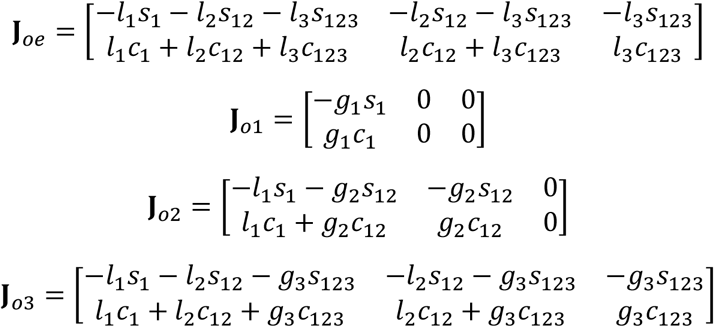

where *l_i_* is the length of link *i* and *g_i_* is the distance from joint *i* − 1 to the center of mass of link *i (estimated from participants’ weight and height* (63)). The notation s_ijk_ and c_ijk_ denotes the functions sin(q_i_ + q_j_ + q_k_) and cos(q_i_ + q_j_ + q_k_), respectively.

## Inverse Kinematics

The inverse kinematics of the 3-link manipulator for a given angular position *q*_1_ of the hip joint is given by (64):

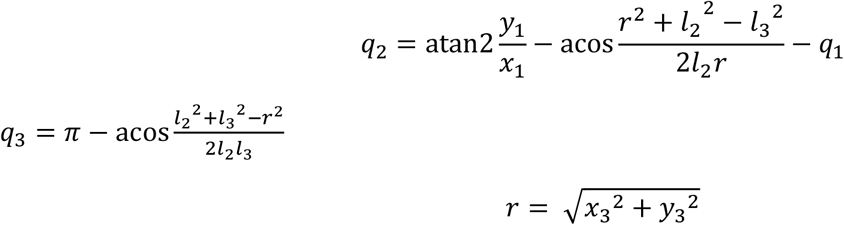

where *x*_3_ and *y*_3_ are the horizontal and vertical components of the position of the end-point with respect to the shoulder joint, and *l*_2_ and *l*_3_ are the lengths of the upper arm and forearm segments, respectively.

## DATA AVAILABILITY

The code and the datasets generated and analyzed during the current study are available in the Open Science Framework repository, https://osf.io/2f3wq/.

## SUPPLEMENTAL MATERIAL

Additional methods and figures are provided in the supplementary material, accessible via the following link: https://osf.io/47szw.

## ACKNOWLEDGMENTS

We would like to thank Simon Pla for helping setup the experiment. Preprint is available at https://doi.org/10.1101/2024.03.28.587142.

## AUTHOR CONTRIBUTIONS

Conceptualization: GF, VB, DM, NS. Data curation: GF, VB. Formal analysis: GF, VB. Funding Acquisition: DM. Investigation: GF. Methodology: GF, VB, DM, NS. Project administration: DM, NS. Resources: DM. Software: GF, VB. Supervision: DM, NS. Validation: GF, VB. Visualization: GF. Writing - Original Draft: GF, VB. Writing - Review & Editing: GF, VB, DM, NS.

## REFERENCES

1. Bernstein. The co-ordination and regulation of movements. Oxford: Pergamon Press, 1967.

2. Todorov E, Jordan MI. Optimal feedback control as a theory of motor coordination. Nature Neuroscience 5: 1226–1235, 2002. doi: 10.1038/nn963.

3. Berret B, Chiovetto E, Nori F, Pozzo T. Evidence for Composite Cost Functions in Arm Movement Planning: An Inverse Optimal Control Approach. PLOS Computational Biology 7: e1002183, 2011. doi: 10.1371/journal.pcbi.1002183.

4. Morel P, Ulbrich P, Gail A. What makes a reach movement effortful? Physical effort discounting supports common minimization principles in decision making and motor control. PLoS Biol 15: e2001323, 2017. doi: 10.1371/journal.pbio.2001323.

5. Prilutsky BI, Zatsiorsky VM. Optimization-Based Models of Muscle Coordination. Exerc Sport Sci Rev 30: 32, 2002.

6. Scott SH. Optimal feedback control and the neural basis of volitional motor control. Nat Rev Neurosci 5: 532–545, 2004. doi: 10.1038/nrn1427.

7. Shadmehr R, Huang HJ, Ahmed AA. A Representation of Effort in Decision-Making and Motor Control. Current Biology 26: 1929–1934, 2016. doi: 10.1016/j.cub.2016.05.065.

8. Henneman E, Somjen G, Carpenter DO. Excitability and inhibitibility of motoneurons of different sizes. Journal of Neurophysiology 28: 599–620, 1965. doi: 10.1152/jn.1965.28.3.599.

9. Milner-Brown HS, Stein RB, Yemm R. Changes in firing rate of human motor units during linearly changing voluntary contractions. J Physiol 230: 371–390, 1973.

10. Milner-Brown HS, Stein RB, Yemm R. The contractile properties of human motor units during voluntary isometric contractions. J Physiol 228: 285–306, 1973. doi: 10.1113/jphysiol.1973.sp010087.

11. Diedrichsen J, Shadmehr R, Ivry RB. The coordination of movement: optimal feedback control and beyond. Trends in Cognitive Sciences 14: 31–39, 2010. doi: 10.1016/j.tics.2009.11.004.

12. McDonald KA, Cusumano JP, Hieronymi A, Rubenson J. Humans trade off whole-body energy cost to avoid overburdening muscles while walking. Proceedings of the Royal Society B: Biological Sciences 289: 20221189, 2022. doi: 10.1098/rspb.2022.1189.

13. Guigon E, Baraduc P, Desmurget M. Computational Motor Control: Redundancy and Invariance. Journal of Neurophysiology 97: 331–347, 2007. doi: 10.1152/jn.00290.2006.

14. Uno Y, Kawato M, Suzuki R. Formation and control of optimal trajectory in human multijoint arm movement. Biological Cybernetics 61, 1989. doi: 10.1007/BF00204593.

15. Cirstea MC, Levin MF. Compensatory strategies for reaching in stroke. Brain 123: 940–953, 2000. doi: 10.1093/brain/123.5.940.

16. Jones TA. Motor compensation and its effects on neural reorganization after stroke. Nat Rev Neurosci 18: 267–280, 2017. doi: 10.1038/nrn.2017.26.

17. Fuller JR, Fung J, Côté JN. Posture-movement responses to stance perturbations and upper limb fatigue during a repetitive pointing task. Human Movement Science 32: 618–632, 2013. doi: 10.1016/j.humov.2013.03.002.

18. Faity G, Mottet D, Pla S, Froger J. The reserve of joint torque determines movement coordination. Sci Rep 11: 23008, 2021. doi: 10.1038/s41598-021-02338-4.

19. Bakhti KKA, Mottet D, Schweighofer N, Froger J, Laffont I. Proximal arm non-use when reaching after a stroke. Neuroscience Letters 657: 91–96, 2017. doi: 10.1016/j.neulet.2017.07.055.

20. Hadjiosif AM, Branscheidt M, Anaya MA, Runnalls KD, Keller J, Bastian AJ, Celnik PA, Krakauer JW. Dissociation between abnormal motor synergies and impaired reaching dexterity after stroke. Journal of Neurophysiology 127: 856–868, 2022. doi: 10.1152/jn.00447.2021.

21. Levin MF, Liebermann DG, Parmet Y, Berman S. Compensatory Versus Noncompensatory Shoulder Movements Used for Reaching in Stroke. Neurorehabil Neural Repair 30: 635–646, 2016. doi: 10.1177/1545968315613863.

22. Nguyen H, Phan T, Shadmehr R, Lee SW. Choice of Arm Use in Stroke Survivors is Largely Driven by the Energetic Cost of the Movement..

23. Taub E, Uswatte G, Elbert T. New treatments in neurorehabiliation founded on basic research. Nature Reviews Neuroscience 3: 228–236, 2002. doi: 10.1038/nrn754.

24. Taylor JL, Gandevia SC. A comparison of central aspects of fatigue in submaximal and maximal voluntary contractions. Journal of Applied Physiology 104: 542–550, 2008. doi: 10.1152/japplphysiol.01053.2007.

25. Enoka RM, Duchateau J. Muscle fatigue: what, why and how it influences muscle function. J Physiol 586: 11–23, 2008. doi: 10.1113/jphysiol.2007.139477.

26. Gandevia SC. Spinal and Supraspinal Factors in Human Muscle Fatigue. Physiological Reviews 81: 1725–1789, 2001. doi: 10.1152/physrev.2001.81.4.1725.

27. Akagi R, Takai Y, Ohta M, Kanehisa H, Kawakami Y, Fukunaga T. Muscle volume compared to cross-sectional area is more appropriate for evaluating muscle strength in young and elderly individuals. Age Ageing 38: 564–569, 2009. doi: 10.1093/ageing/afp122.

28. Fukunaga T, Miyatani M, Tachi M, Kouzaki M, Kawakami Y, Kanehisa H. Muscle volume is a major determinant of joint torque in humans. Acta Physiol Scand 172: 249–255, 2001. doi: 10.1046/j.1365-201x.2001.00867.x.

29. Maughan RJ, Watson JS, Weir J. Strength and cross-sectional area of human skeletal muscle. J Physiol 338: 37–49, 1983. doi: 10.1113/jphysiol.1983.sp014658.

30. Danneskiold-Samsøe B, Bartels EM, Bülow PM, Lund H, Stockmarr A, Holm CC, Wätjen I, Appleyard M, Bliddal H. Isokinetic and isometric muscle strength in a healthy population with special reference to age and gender. Acta Physiologica 197: 1–68, 2009. doi: 10.1111/j.1748-1716.2009.02022.x.

31. Borchers HW. pracma: Practical Numerical Math Functions [Online]. 2022. https://CRAN.R-project.org/package=pracma [11 Apr. 2023].

32. Wickham H, Chang W, Henry L, Pedersen TL, Takahashi K, Wilke C, Woo K, Yutani H, Dunnington D, Posit, PBC. ggplot2: Create Elegant Data Visualisations Using the Grammar of Graphics [Online]. 2023. https://CRAN.R-project.org/package=ggplot2 [11 Apr. 2023].

33. Raftery AE. Bayesian Model Selection in Social Research. Sociological Methodology 25: 111–163, 1995. doi: 10.2307/271063.

34. Andrews AW, Bohannon RW. Distribution of muscle strength impairments following stroke. Clin Rehabil 14: 79–87, 2000. doi: 10.1191/026921500673950113.

35. Fuglevand AJ, Winter DA, Patla AE. Models of recruitment and rate coding organization in motor-unit pools. J Neurophysiol 70: 2470–2488, 1993. doi: 10.1152/jn.1993.70.6.2470.

36. Canning CG, Ada L, Adams R, O’Dwyer NJ. Loss of strength contributes more to physical disability after stroke than loss of dexterity. Clin Rehabil 18: 300–308, 2004. doi: 10.1191/0269215504cr715oa.

37. Mercier C, Bourbonnais D. Rlative shoulder fexor and handgrip strength is related to upper limb function after stroke. Clin Rehabil 18: 215–221, 2004. doi: 10.1191/0269215504cr724oa.

38. McCrea PH, Eng JJ, Hodgson AJ. Saturated Muscle Activation Contributes to Compensatory Reaching Strategies After Stroke. Journal of Neurophysiology 94: 2999–3008, 2005. doi: 10.1152/jn.00732.2004.

39. Lieshout ECC van, van de Port IG, Dijkhuizen RM, Visser-Meily JMA. Does upper limb strength play a prominent role in health-related quality of life in stroke patients discharged from inpatient rehabilitation? Topics in Stroke Rehabilitation 27: 525–533, 2020. doi: 10.1080/10749357.2020.1738662.

40. Valdés BA, Glegg SMN, Van der Loos HFM. Trunk Compensation During Bimanual Reaching at Different Heights by Healthy and Hemiparetic Adults. Journal of Motor Behavior 49: 580–592, 2017. doi: 10.1080/00222895.2016.1241748.

41. Krakauer JW, Carmichael ST. Broken Movement: The Neurobiology of Motor Recovery after Stroke. MIT Press, 2022.

42. Carr JH, Shepherd RB. A Motor Learning Model for Stroke Rehabilitation. Physiotherapy 75: 372–380, 1989. doi: 10.1016/S0031-9406(10)62588-6.

43. Kim S, Han CE, Kim B, Winstein CJ, Schweighofer N. Effort, success, and side of lesion determine arm choice in individuals with chronic stroke. Journal of Neurophysiology 127: 255–266, 2022. doi: 10.1152/jn.00532.2020.

44. Latash ML, Anson JG. What are “normal movements” in atypical populations? Behavioral and Brain Sciences 19: 55–68, 1996. doi: 10.1017/S0140525X00041467.

45. Engelbrecht SE. Minimum Principles in Motor Control..

46. Flash T, Hogan N. The coordination of arm movements: an experimentally confirmed mathematical model. J Neurosci 5: 1688–1703, 1985. doi: 10.1523/JNEUROSCI.05-07-01688.1985.

47. Harris CM, Wolpert DM. Signal-dependent noise determines motor planning. Nature 394: 780, 1998. doi: 10.1038/29528.

48. Nelson WL. Physical principles for economies of skilled movements. Biological Cybernetics 46: 135–147, 1983. doi: 10.1007/BF00339982.

49. Todorov E. Optimality principles in sensorimotor control. Nature Neuroscience 7: 907, 2004. doi: 10.1038/nn1309.

50. Alexander RM. Optimization and gaits in the locomotion of vertebrates. Physiological Reviews 69: 1199–1227, 1989. doi: 10.1152/physrev.1989.69.4.1199.

51. Ralston HJ. Energy-speed relation and optimal speed during level walking. Int Z Angew Physiol Einschl Arbeitsphysiol 17: 277–283, 1958. doi: 10.1007/BF00698754.

52. Selinger JC, O’Connor SM, Wong JD, Donelan JM. Humans Can Continuously Optimize Energetic Cost during Walking. Current Biology 25: 2452–2456, 2015. doi: 10.1016/j.cub.2015.08.016.

53. Zarrugh MY, Todd FN, Ralston HJ. Optimization of energy expenditure during level walking. Europ J Appl Physiol 33: 293–306, 1974. doi: 10.1007/BF00430237.

54. Ackermann M, van den Bogert AJ. Optimality principles for model-based prediction of human gait. Journal of Biomechanics 43: 1055–1060, 2010. doi: 10.1016/j.jbiomech.2009.12.012.

55. Miller RH, Umberger BR, Hamill J, Caldwell GE. Evaluation of the minimum energy hypothesis and other potential optimality criteria for human running. Proceedings of the Royal Society B: Biological Sciences 279: 1498–1505, 2011. doi: 10.1098/rspb.2011.2015.

56. Gulletta G, Silva EC e, Erlhagen W, Meulenbroek R, Costa MFP, Bicho E. A Human-like Upper-limb Motion Planner: Generating naturalistic movements for humanoid robots. International Journal of Advanced Robotic Systems 18: 1729881421998585, 2021. doi: 10.1177/1729881421998585.

57. Faity G, Mottet D, Bakhti K, Froger J, Laffont I. Shoulder weakness induces trunk compensation after a stroke [Online]. 16th world congress of the ISPRM https://hal.science/hal-03778164 [11 Apr. 2023].

58. Ada L, Dorsch S, Canning CG. Strengthening interventions increase strength and improve activity after stroke: a systematic review. Australian Journal of Physiotherapy 52: 241–248, 2006. doi: 10.1016/S0004-9514(06)70003-4.

59. Dorsch S, Ada L, Alloggia D. Progressive resistance training increases strength after stroke but this may not carry over to activity: a systematic review. Journal of Physiotherapy 64: 84–90, 2018. doi: 10.1016/j.jphys.2018.02.012.

60. Harris Jocelyn E., Eng Janice J. Strength Training Improves Upper-Limb Function in Individuals With Stroke. Stroke 41: 136–140, 2010. doi: 10.1161/STROKEAHA.109.567438.

61. Lum PS, Mulroy S, Amdur RL, Requejo P, Prilutsky BI, Dromerick AW. Gains in Upper Extremity Function After Stroke via Recovery or Compensation: Potential Differential Effects on Amount of Real-World Limb Use. Topics in Stroke Rehabilitation 16: 237–253, 2009. doi: 10.1310/tsr1604-237.

62. Georgarakis A-M, Xiloyannis M, Wolf P, Riener R. A textile exomuscle that assists the shoulder during functional movements for everyday life. Nat Mach Intell 4: 574–582, 2022. doi: 10.1038/s42256-022-00495-3.

63. De Leva P. Adjustments to zatsiorsky-seluyanov’s segment inertia parameters. Journal of Biomechanics: 1996.

64. Murray RM, Li Z, Shankar Sastry S. A mathematical introduction to robotic manipulation. 2017.

