## Supplementary Material for "Force Reserve Predicts Compensation in Reaching Movement with Induced Shoulder Strength Deficit" (from another version of this preprint: version 1, 2024-03-30)

RESEARCH ARTICLE



### Supplementary material

*Methods - Joint Torque Computation*

In our seated reaching task, participants performed movements that approached static conditions. Specifically, they were instructed to touch a target located in front of them, maintain their hand at the target for a duration of 1 second, and then return their arm to the armrest of the chair. Given the quasi-static nature of this task, the primary determinants of torque at various joints arise predominantly from gravitational forces. This context allows us to adopt a static analysis approach rather than a comprehensive inverse dynamics analysis, ensuring both computational efficiency and precision appropriate for our experimental conditions.

Therefore, the torque around a given joint is determined by the sum of the torques generated by the weights of the load and the body segments that the joint supports

$$\boldsymbol{\tau}_i = \boldsymbol{\tau}_{L,i} + \sum_{j=1}^{3} \boldsymbol{\tau}_{w,ij} \qquad (supp.1)$$

where $\boldsymbol{\tau}_i$ is the 3D static torque around joint $i$ = 1, 2, 3 (hips, shoulder, elbow), $\boldsymbol{\tau}_{L,i}$ is the torque generated by the dumbbell around joint $i$, and $\boldsymbol{\tau}_{w,ij}$ is the torque generated by the weight of body segment $j$ = 1, 2, 3 (trunk, upper arm, and forearm-hand) around joint $i$. Note that $\boldsymbol{\tau}_{w,ij}$ = 0 for $i > j$, as this indicates a torque from body segment $j$ that is not supported by joint $i$ (i.e. $\boldsymbol{\tau}_{w,31}$ = 0). We calculated the gravitational torques $\boldsymbol{\tau}_{w,ij}$ as the cross product of the lever arm of body segment $j$ to joint $i$ and the weight vector of the body segment $j$:

$$\boldsymbol{\tau}_{w,ij} = (\boldsymbol{r}_{COM,j} - \boldsymbol{r}_i) \times \boldsymbol{w}_j \qquad (supp.2)$$

where $\boldsymbol{r}_{COM,j}$ and $\boldsymbol{r}_i$ are position vectors of the center of mass of body segment $j$ and joint $i$, respectively, with respect to the ground frame, and $\boldsymbol{w}_j$ is the weight vector of body segment $j$ ($\boldsymbol{w}_L = [0\ 0\ -w_L]^T$, where $w_L$ is the absolute weight of the body segment). The position of joints $\boldsymbol{r}_i$ was derived from motion capture measurements. Positions of the center of mass and the absolute weight of the body segments were derived from participants' height and weight using De Leva's equations (57). We calculated the load torques $\boldsymbol{\tau}_{L,i}$ in a similar way.

We defined the antigravity joint torques as the Euclidean norm of the 3D static torques $\boldsymbol{\tau}_i$ at the terminal posture at $t_{\text{final}}$. For each participant, we normalized the antigravity joint torques with respect to the participant's maximum voluntary torque (MVT) at the corresponding joint, such that the torque at the joint is expressed as a percentage of the MVT.

*Results - Adaptive Strategies and Leg Use*

Our experimental data also captured various adaptive strategies when the arm's weight increased. These strategies encompassed enhanced leg recruitment, trunk rotation, and shoulder abduction as compensatory movements, as illustrated in Figure Supp. 1. Analysis of hip displacement during reaching movements pinpointed the initiation of leg recruitment at approximately 50-60% shMVT. This onset is concomitant with a reduced shoulder force reserve. Specifically, for a dumbbell weight of ≤ 10% shMVT, we observed a hip displacement of 0.4% ±1.1 of the reach length, trunk rotation of 13.1° ±7.2, and shoulder abduction of 7.2° ±8.2. In contrast, for dumbbell weights ranging from 40 to 60% shMVT, the respective values were 2.0% ±1.2 for hip displacement, 21.0° ±6.7 for trunk rotation, and 13.3° ±12.1 for shoulder abduction (Figure Supp. 1A). Upon surpassing the aforementioned threshold, exclusive reliance on trunk movement proved insufficient for conserving force reserve, prompting the adoption of additional compensatory movement strategies. In situations where participants' shoulder torque exceeded the 60-70% MVT threshold, a subtle disengagement of the pelvis from the chair was observed, manifesting as a simple rocking motion. For dumbbell weights between 90-110% shMVT, the measurements were 12.1% ±6.4 for hip displacement, 21.9° ±7.5 for trunk rotation, and 21.5° ±14.1 for shoulder abduction. When dumbbell weights ranged from 140-160% shMVT, the corresponding values were 20.8% ±4.2 for hip displacement, 15.2° ±10.1 for trunk rotation, and 35.1° ±10.4 for shoulder abduction. In the heaviest dumbbell conditions, particularly when arm weights surpassed 150% shMVT (equating to 15 and 18 kg), a subset of participants opted to standing to accomplish the reaching task. This adjustment was observed in participants with the lowest maximum shoulder torque capabilities (Figure Supp. 1B).

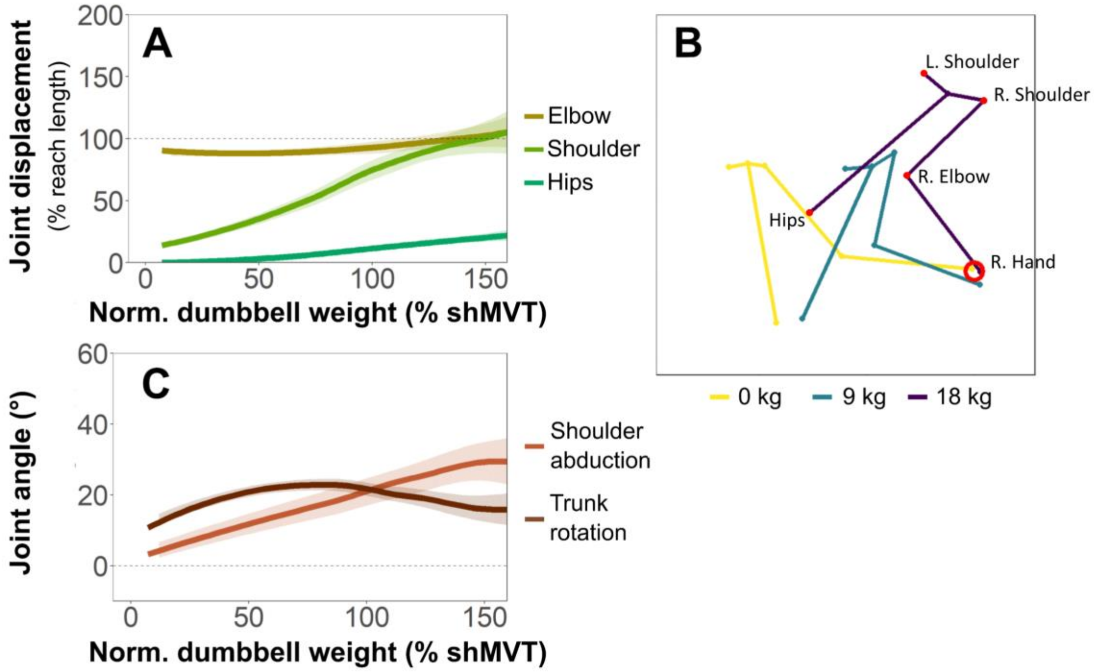


***Supplementary data #1****. Model Limitations vs. Experimental Data Complexity. Panel A: Elbow, shoulder, and hip displacements during the reach transition from the initial resting posture to the final posture, plotted against normalized dumbbell weight. An escalation in dumbbell weight accentuates trunk involvement, evident from the increased displacements of the shoulder joint. Moreover, leg involvement initiates at 50% shMVT, marked by heightened displacement of the hips. For panel A and panel C, as detailed in the Methods section, the X-axis represents the normalized dumbbell weight as a percentage of the participants' shoulder maximum voluntary torque (% shMVT). Thick lines show the overall average, whereas the shaded areas indicate the 95% confidence interval of the mean. Panel B: 2D projection of postures from a representative subject plotted against dumbbell weight. The shifting base of the trunk highlights the subject's strategy of incrementally moving their hips closer to the target in tandem with rising dumbbell weight, culminating in the subject rising from their chair at extreme weights (>150% shMVT). Panel C: Plotted angles of shoulder abduction (light brown) and trunk rotation (dark brown) against normalized dumbbell weight. Shoulder abduction witnesses an initial surge up to approximately 70% shMVT, followed by a decline in the subsequent phase. In contrast, trunk rotation steadily amplifies with increasing dumbbell weight.*
